# Beyond community-weighted means: quantifying trait distributions for detecting community assembly patterns

**DOI:** 10.1101/2024.12.18.629105

**Authors:** Yolanda Melero, Constanti Stefanescu, David García-Callejas

## Abstract

The distributions of ecological traits are commonly used to infer the processess structuring ecological communities, as these processes (either deterministic or stochastic) select and filter species, leading to distint trait patterns across communities. Trait distributions are frequently characterised by their average, the community-weighted mean, but increasing evidence exists for non-gaussian trait distributions in empirical communities. In such situations, community-weighted means are insufficient to capture the patterns of community traits and to infer the implied ecological processes. Here we analyse the empirical distributions of 6 functional traits of butterflies from a natural community and a filtered community from a urban area across a period of six years. First, we show that to adequately describe trait distributions, statistical descriptors beyond community-weighted means are needed, as distributions were in all cases clearly non-gaussian. In particular, besides distribution averages and standard deviations, we compute the skewness, kurtosis, the range of trait values, and a multimodality index. Second, we compare this set of descriptors between our natural and filtered communities. We find clear differences between communities, detected in particular by combining the relationships between the skewness, kurtosis, and range of the distributions. These analyses allow us to infer that the filtered community is mostly shaped by deterministic filtering processes, through a mixture of directional and stabilizing assembly filters. These patterns are furthermore consistent across the six years of data, providing further evidence of deterministic processes shaping the assembly of our urban butterfly community. Overall, we provide evidence of the ubiquity of non-gaussian trait distributions across natural and filtered communities, propose key descriptors for understanding and comparing such distributions, and identify filtering process structuring these communities.

## Introduction

Ecological traits, the measurable features (morphological, ecological or behavioural) of an organism (Calow, 1987; Dawson *et al*., 2021), underpin the relationships between individuals, populations and communities with their environment (de Bello *et al*., 2021). Concurrently, environmental and biotic constraints dictate the individuals that thrive in a given context (population, community, ecosystem), and thus the observed values of a specific trait are the result of several deterministic and stochastic ecological and evolutionary processes (Chase & Myers, 2011; Enquist *et al*., 2015; Macarthur & Levins, 1967). For example, at the population level, the colonization of a new site is strongly dependent on both stochastic and deterministic dispersal processes, such as random arrival of individuals or density-dependent dispersal. Trait values of the individuals at the colonised site will be a subset of those from the source sites, mediated by the relative importance of the different dispersal processes (Clobert *et al*., 2012). The set of values of a trait that are observed in a given assemblage then define the frequency distribution of that trait (Enquist *et al*., 2015).

Trait distributions are generally defined at the species (Bolnick *et al*., 2011, 2003) or more commonly at the community level (McGill *et al*., 2006). In animal assemblages, these distributions are often analysed by summarising them using weighted means. When analysing ecological communities, the commonly-used Community-Weighted Mean (CWM) is defined as the mean value of the trait among all species in the studied communities weighted by their abundances, which is then used to compare different communities, in itself or as part of other analyses such as the fourth-corner approach or other correlation methods (Miller *et al*., 2019; Peres-Neto *et al*., 2017). Less frequently, the variance or the standard deviation of the CWM is also provided, as an indicator of the variability of the trait at the level of organisation studied (Mitchell *et al*., 2021). Implicit in the use of these metrics (CWM and its variance or standard deviation) is the assumption that trait distributions follow a gaussian shape, and therefore the CWM represents the most frequent trait value, with other values symmetrically declining in frequency. In ecological terms, gaussian distributions assume that there is a unique trait value that best matches the abiotic and biotic environment under study, i.e. a value with the highest fitness, deviations from which imply a symmetrical reduction in fitness. Trait distributions, however, are not necessarily gaussian (e.g. animal body size is frequently skwewed (Koz-lowski & Gawelczyk, 2002)). Further, maximum fitness values may be achieved at different values of a given trait (multimodality) with suboptimal trait values below these maxima. For example, pollination effectiveness in bee communities can be maximized via both large or medium body size (Roquer-Beni *et al*., 2022); likewise, flight performance in butterfly communities is dominated by both small (high maneuverability) and large wing sizes (high flight duration) (Le Roy *et al*., 2019).

These simplifications of community trait distributions into Community-Weighted Means and, eventually, their standard deviation, thus leave out key information about non-gaussian trait distributions, including its potential asymmetry, evenness, and multimodality. In statistical distributions, the mean represents their first moment, and the higher moments are the variance or standard variation, the skewness (which measures the asymmetry of the distribution) and kurtosis (which informs about the peaks and tails of the distribution). Theoretical and empirical studies have shown that in order to infer community assembly processes, a more complete characterisation of trait distributions is necessary, including the relationship between the moments of the distributions (Denny, 2017; Gross *et al*., 2017, 2021). In tree communities across the globe, enviromental factors have been shown to modify not only mean trait values, but also standard deviation, skewness and kurtosis (Wieczynski *et al*., 2019). Likewise, skewness and kurtosis are important to understand changes in body size distributions after species invasions (e.g. in freshwater fish communities; (Blanchet *et al*., 2010)), metrics that can vary independently of the mean and variance specially in response to environmental gradients (e.g. in plants; (Le Bagousse-Pinguet *et al*., 2017)). These considerations are especially important for understanding differences between communities with potentially similar species pools, but in which environmental and/or dispersal filters are known to operate. This is because these filters are known to select trait values in non-uniform ways from the source communities, altering not only the mean but also the range and higher moments of the source distributions (Perronne *et al*., 2017). Current frameworks for understanding trait-based filtering processes are however based on gaussian trait distributions (Lavorel *et al*., 2007; Lep̌s & de Bello, 2023), and their applicability in more complex situations remains unclear.

Urban communities are an ideal setting to test the effects of filters in trait distributions and the limitations of Community-Weighted Means at capturing these effects. Urban environments present strong filters because for many species they represent novel conditions to which they are not adapted, hence being absent or rare. Species whose traits confer them high adaptability may in turn dominate the urban community (Callaghan *et al*., 2021; Santini *et al*., 2018). In these environments, changes in mean trait values in response to urbanisation have been documented for plant (Palma *et al*., 2017) and terrestrial animal species worlwide (Hahs *et al*., 2023). However, there is little evidence on how urban filters influence the range, multimodality, or the higher moments of trait distributions compared to their source communities. We expect that, in general, the range of trait values observed in urban communities will be reduced compared to their source. The resulting trait distributions in urban communities may also reflect environmental biases associated to increased peaks at optimum trait values and lower tails (Enquist *et al*., 2015; Gross *et al*., 2017; Keddy, 1992), or similar trait distribution shapes if the frequency of traits is the result of random dispersal from the source community.

The consistency of trait distributions with time is, likewise, key to understand ecological processes of community assembly and dynamics, particularly in filtered communities. If trait distributions are conserved across time, this may imply deterministic dispersal events from the regional source, and/or ecological processes modulating the intrinsic dynamics of urban communities. In turn, a high variability in trait distributions of filtered communities may be indicative of stochastic dispersal or local factors driving community structure and dynamics of these communities (Grime, 2006; Norberg *et al*., 2001).

Here we studied spatio-temporal changes in the distribution of six representative traits of two butterfly communities across six years. We analyzed a regional (source) community in comparison with a filtered urban assemblage (henceforth we use “community” to refer to these broad assemblages, for simplicity). We used butterflies because they show a broad and well studied range and variation in life-history and ecological traits (Middleton-Welling *et al*., 2020), and respond quickly to environmental change minimizing the effect of time-lag responses (Devictor *et al*., 2012; Krauss *et al*., 2010). Moreover, they are representative of other animal species, ensuring reliable general ecological patterns (Thomas, 2005). We characterised the four moments of the trait distributions, their range and multimodality, and used these metrics to evaluate potential processes driving community assembly in the urban environment.

## Methods

### Species traits and data collection

We conducted our study in the province of Barcelona (NE Spain). The regional community included the (semi)natural areas surrounding the municipality of Barcelona up to 60km, sharing the same bioclimatic conditions and altitudinal levels as those of the urban area (warm, xeric climate and *<* 600m of altitude; Metzger *et al*. (2013). The filtered urban community (hencefort “filtered”) includes butterfly species observed in urban parks within the city of Barcelona. We monitored 129 sites for the regional community and 29 parks within the filtered community. Sampling was carried out by volunteers involved in two citizen science programs: the “Catalan Butterfly Monitoring Scheme” (CBMS, www.catalanbms.org) and the “urban Butterfly Monitoring Scheme” (uBMS, www.ubms.creaf.cat). Individuals were counted weekly from March to September following the international-standard “Pollard walk” methodology Pollard (1977) along fixed transects (CBMS) and along additional random walks of fixed temporal duration to account for small isolated sampling sites within the municipality (uBMS). We used data collected between 2018 and 2023. This led to a total of 589,316 and 22,551 observations from 160 and 54 morphospecies in the CBMS and the uBMS respectively (further details of the sampling protocol and species definition in Methods S1).

We calculated the annual abundance of adult butterflies for each species and community *N_ijt_*where *i* is the species, *j* the community (regional or filtered) and *t* is the year. Annual abundances were calculated from the raw count data using the standardized annual abundance index (Schmucki *et al*., 2016). This index consists first of a generalized additive model (GAM) with a Poisson distribution and log link function that calculates the phenology (i.e. the adult flight curve) of a species per community and year; and second, it uses a generalized log-linear model with the species phenology as offset to predict missing counts and estimate the species annual abundance index per community and year (Dennis *et al*., 2016; Schmucki *et al*., 2022, 2016). This method is considered more robust than linear interpolation since it takes in account species phenology, and it is widely used to obtain butterfly species abundances with the European Butterfly Monitoring Scheme (Montgomery *et al*., 2021; Van Swaay *et al*., 2022). To reduce inconsistencies we excluded indices of species containing *>* 50% of missing data per community. To avoid losing species and traits at the tails of the distribution, we extrapolated the annual abundance indexes of those species excluded due to sporadic observations using linear regression (N_CBMS_ = 76 missing species and year combinations, of 25 different species; N_uBMS_ = 10 missing species and year combinations, of 7 different species; details in Methods S2).

We selected six quantitative traits from butterfly species linked to their ecology, behaviour and morphology (Boggs *et al*., 2003; Dennis, 1992, 2010). Two of the traits selected relate to trophic specialization: (i) the host-plant specialization index (HSI), which quantifies the trophic specialization of a butterfly species in the larval stage based on the number of plant taxa it feed upon, standarized to a range of 0-1 (most generalist to most specialist), and (ii) the adult species specialisation index (SSI), measured as the relative average butterfly species density per CORINE habitat type and standarized to empirically ranging 0-4 (most generalist to most specialist). Two further traits relate to butterfly climatic niche and, indirectly, thermal tolerance: (iii) species temperature index (STI), a proxy for the species climatic niche centroid set as the mean temperature within the species distributional range; and (iv) species temperature variation index (SVTI), a proxy for the species climatic niche breadth set as the standard deviation of the temperature within the species range. We also included one trait related to habitat use: (v) the degree of preference of closed to open areas (TAO index) calculated as the relative butterfly density at open versus closed pre-categorized areas, ranging −1 to 1. Finally, we used one trait related to body size: (vi) wing index (WI), relating both the forewing length and the wingspan of both females and males. Trait values per species were extracted from Middleton-Welling *et al*. (2020); Schweiger *et al*. (2014) and Ubach *et al*. (2020). Due to data limitations, trait values are aggregated at the species level, without considering intra-specific variability (but see below for bootstrap estimations of intra-specific variation).

### Trait distribution descriptors

We assessed the abundance-weighted distributions of the six species traits per community and calculated the four moments defining each distribution: weighted mean (i.e. the CWM), standard deviation, skewness and kurtosis. We further obtained the range of each trait distribution, defined as the absolute value of the difference between the maximum and minimum values of each distribution, and the unior multimodality of each species distribution. In addition, we calculated the relationship between kurtosis and skewness, defined as (*K ≥ β · S*^2^ + *α*). This relationship defines a theoretical lower boundary of kurtosis depending on skewness, that in the context of trait distributions represents the maximum value of trait diversity for a given skewness value. There-fore, the observed *K − S*^2^ relationship from a given empirical distribution informs about the trait diversity of the distribution in relation to that potential maximum (Gross *et al*., 2017, 2021). The *K − S*^2^ relationship in a gaussian distribution is zero, as kurtosis equals three but skewness is zero: departures from these values are therefore used to characterize non-gaussian distributions.

We calculated the four moments of our trait distributions using the traitstrap R package v0.1.0 (Maitner *et al*., 2023). The bootstrap method implemented in that package for obtaining community-level distribution metrics allows statistically estimating intra-specific trait variability in the absence of further information, thus resulting in distribution metrics with associated confidence intervals (Maitner *et al*., 2023). We used the Hartigan’s Dip test for assessing the potential departure of unimodality in our distributions (Hartigan & Hartigan, 1985), as implemented in the diptest R package v0.77-1 (Maechler, 2024). Lastly, we calculated the coefficient of variation of each metric and trait across years to assess the temporal variabily of each community.

## Results

The distributions of the six traits studied were highly heterogeneous, with none of them showing a gaussian shape, as exemplified in the distribution of two traits in 2019 (Fig. 1 and Fig. S2). Most of the distributions were asymmetrical and showed relatively long tails, as evidenced by high values of skewness and kurtosis (Fig. 2). Moreover, the regional and filtered communities showed consistent differences in most trait distributions. In particular, average trait values (the CWM) tended to be higher for the regional community, except for SVTI (the proxy of climatic niche breadth) and WI (the proxy of body size). Likewise, standard deviation was almost always higher in the regional distributions, except again for WI. The range of trait values observed was also higher, for all traits and years, in the regional communities except for HSI (the host-plant specialization index), which showed the same range in the regional and filtered communities. Lastly, the DIP test of multimodality indicated statistically significant departure from unimodality in all cases. The DIP index was consistently higher in the filtered communities, indicating the co-dominance of different functional groups, if all with lower abundances than the dominant values at the regional community. These dominant values did not coincide with the CWM for any trait (e.g. Fig 1).

**Fig. 1:**
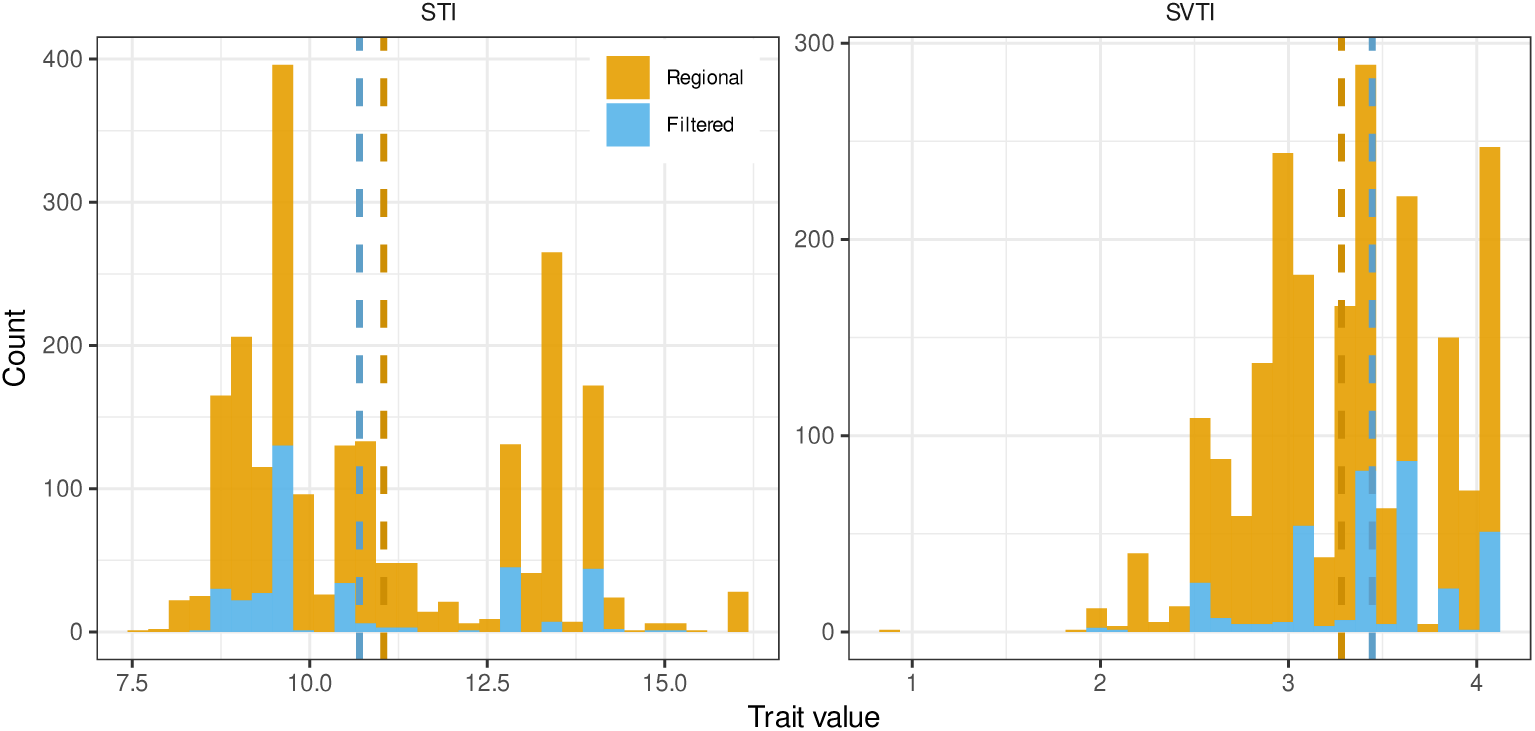
Distributions of STI, SVTI in 2019. Dashed lines indicate the means.

**Fig. 2:**
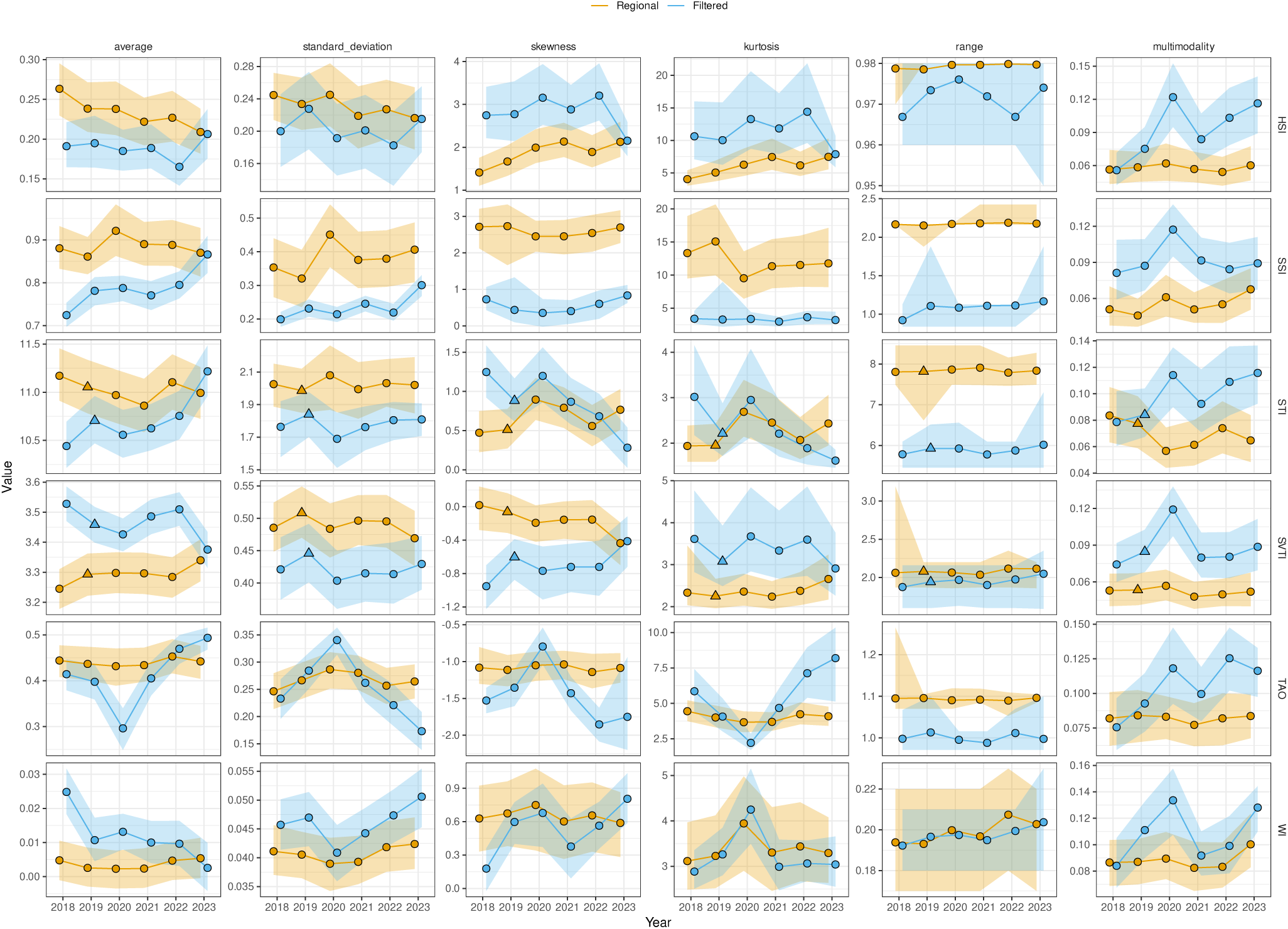
Values of trait distribution metrics across years. 95% confidence intervals are obtained by bootstrapping the raw distributions. Range is simply defined as the absolute value of the difference between the minimum and maximum values observed. Multimodality is calculated with the Hardigan’s DIP test, whereby higher values indicate stronger departure from unimodality. Triangle shapes in 2019 for STI and SVTI are used to highlight the distributions shown in Fig. 1.

The skewness-kurtosis relationships (*S*^2^ *− K*) allowed clearly differentiating the regional and filtered distributions in all traits except WI (Fig. 3). Interestingly, trait diversity of the filtered communities according to the *S*^2^ *− K* relationship was higher for SSI, STI, and WI (lower panels of Fig. 3). This result, however, needs to be understood in parallel with the range of values of each distribution: the range of values of all traits (except WI) was higher in the regional communities (Fig. 2), so kurtosis and skewness values, and their derived trait diversity measurements are not comparable between distributions without considering differences in their ranges, when one range is potentially a subset of the other.

**Fig. 3:**
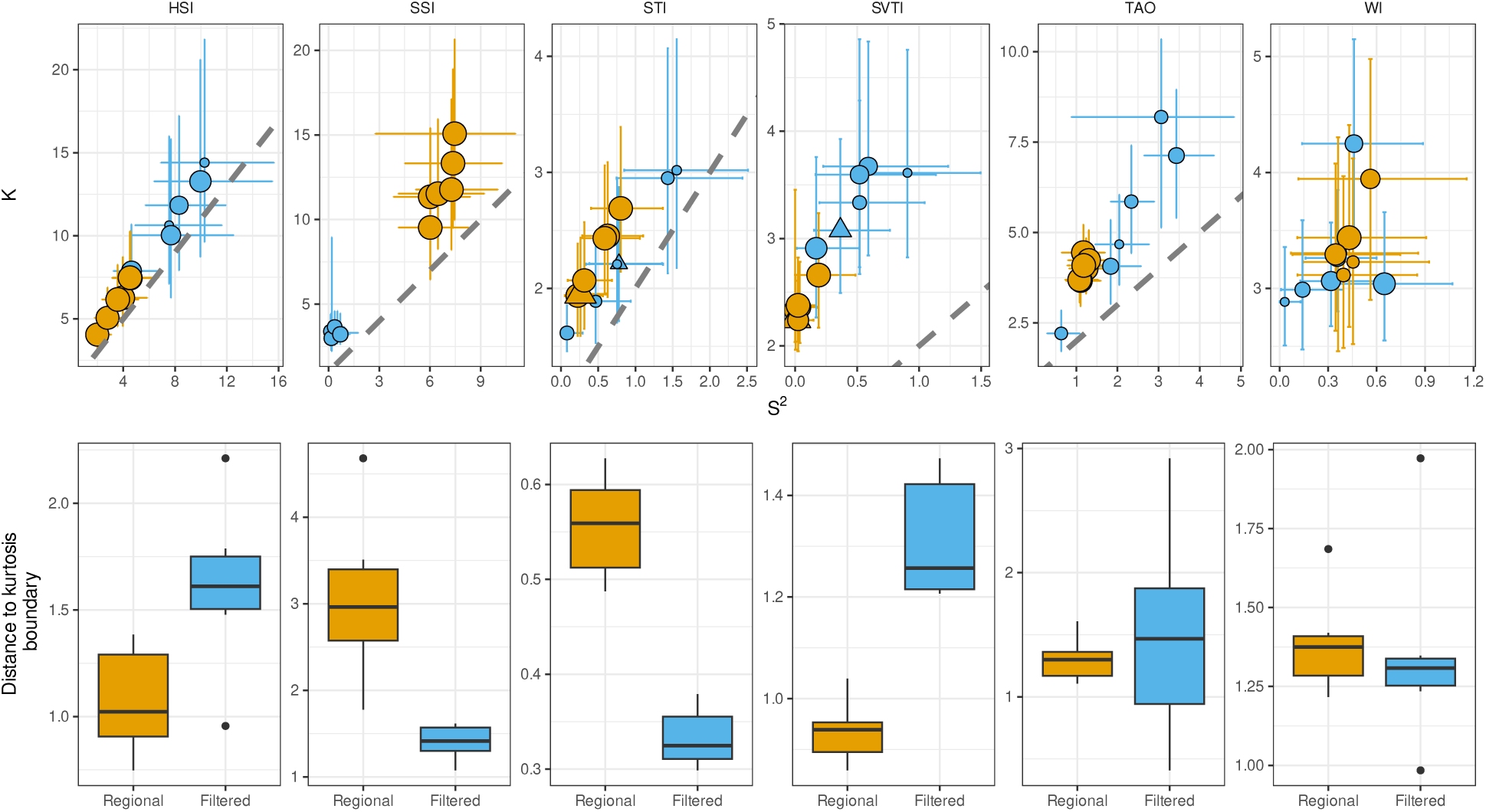
Skewness-kurtosis relationships in the trait distributions studied. Upper panels represent the position of each distribution in the plane defined by kurtosis and skewness squared. Point size indicates the relative range of the distributions, with larger points representing larger average ranges (see Fig 2 for the actual range values). Vertical and horizontal lines represent 95% confidence intervals. The dashed grey lines are the boundary given by *S*^2^ + 1 = *K*, which indicates the minimum value of kurtosis for a given skewness. Triangle shapes for STI and SVTI are used to highlight the distributions shown in Fig. 1. Lower panels represent the distributions of distances from each distribution to that theoretical boundary. In lower panels, the horizontal black line represents the median, the upper and lower hinges correspond to the 25th and 75th percentiles, and the vertical lines extend to the largest/smallest value up to 1.5 times the interquartile range (distance between 25th and 75th percentiles). Single points are observations outside of 1.5 times the interquantile range.

**Table 1:** Descriptors of the distributions from Fig 1.

|  | STI |  | SVTI |  |
| --- | --- | --- | --- | --- |
|  | Regional | Filtered | Regional | Filtered |
| average | 11.04 | 10.69 | 3.28 | 3.44 |
| st. dev. | 1.98 | 1.84 | 0.51 | 0.45 |
| skewness | 0.52 | 0.88 | -0.1 | -0.59 |
| kurtosis | 1.9 | 2.21 | 2.42 | 3.06 |
| range | 8.45 | 6.6 | 3.19 | 2.16 |
| multimodality | 0.07 | 0.08 | 0.05 | 0.08 |

Trait distributions in both the regional and filtered communities were consistent across years (Fig. 4). In particular, the coefficient of variation of 61 out of 72 values was *<* 0.25, and a further 9 values were *<* 0.5. No trait showed a higher variability across all metrics than others: for example, the Wing Index was more variable in its average values across years than other traits, but this was not the case for standard deviation or the other metrics. Notwithstanding these overall trends, the filtered community showed comparatively larger temporal variability than the regional one, with higher coefficient of variation in 30 out of 36 pairs of values, especially for the skweness, kurtosis, and multimodality (Fig. 4).

**Fig. 4:**
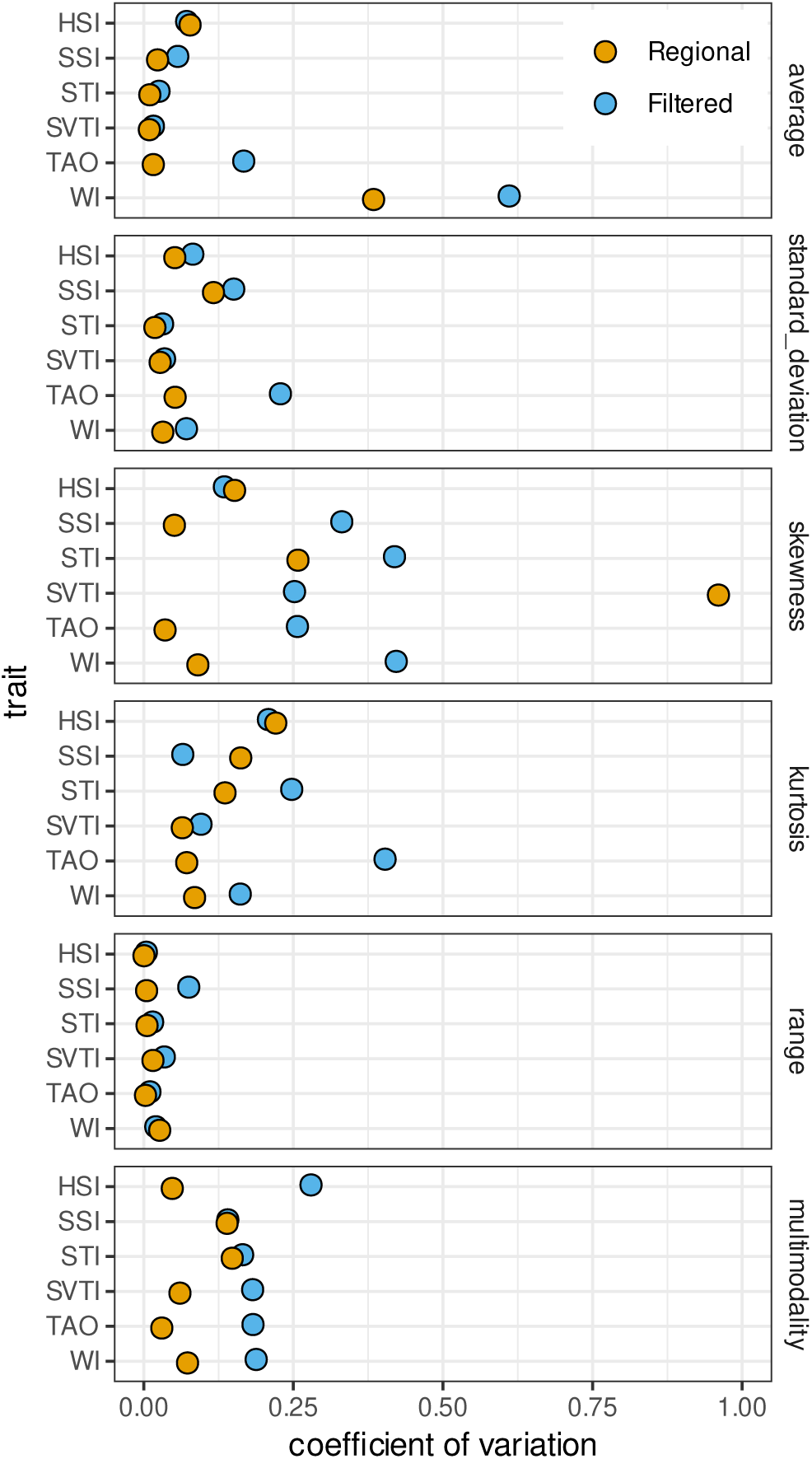
Coefficient of variation of distribution metrics for each trait across years, for the regional and filtered communities. A high CV indicates large temporal variability of that metric, whereas a CV closer to 0 indicates a relatively constant value across years.

### Trophic generalism: larval and adult specialisation index

Species in the regional and filtered communities varied in relation to their trophic specialisation. In general, the regional community was composed of more adult specialist species than the filtered community as indicated by the higher average and standard deviation in SSI (Fig. 2). Potential differences in the larval specialisation (HSI) were partly observable in higher average values of the regional community, although with overlapping confidence intervals. However, both communities showed a dominance of trophic generalist species (right-skewed distributions and high kurtosis), especially in the larval stage of the filtered community. Likewise, there were no extreme adult specialists in the filtered community, as showed by the reduced range in SSI (Fig. 2 and Fig. S2).

The *S*^2^ *− K* relationship showed a clear distinction between both communities (upper panels of Fig. 3); a higher trait diversity in the regional community for larval specialisation, and in the filtered community for adult specialisation (lower panels of Fig. 3). However, the higher diversity of adult specialisation in the filtered community needs to be understood within the constrained range of values (markedly higher in the regional community; Fig. 2).

### Niche position and thermal tolerance

Species from both communties also varied in relation to their climatic niche (STI and STVI), with the filtered community dominated by species with colder centroids (lower STI) and wider climatic niche breadth (higher STVI), i.e. thermal generalist species than the regional community (as shown by their CWMs).Moreover, STVI was more symetrical (lower skewness) and showed higher co-dominances (multimodality) than the regional community, excluding species with cold centroids and narrow climatic niches (Fig. 2 and Fig. S2).

Similarly to the trophic specialisation indices, the *S*^2^ *−K* relationship allowed further insights in the differences between the two communities. The regional community displayed greater distances to the kurtosis boundary in STI, indicating lower trait diversity (lower panels of Fig. 3), which again is countered by a significantly higher range of trait values. For STVI, the regional community showed both higher range values and higher trait diversity than the filtered community.

### Affinity for open areas

Both communties had similar distribution in relation to the species affinity for open areas (TAO), but the filtered community showed higher temporal variability in all metrics except in its reduced range (lacked of species preferring very open areas). The filtered community also had higher frequency of co-dominant species, specially determined by higher (and more variable) appearence of species of closed areas (as seen in its kurtosis, skeweness and multimodality; Fig. 2 and Fig. S2).

As with previous traits, the regional community showed lower trait diversisty (*S*^2^ *− K*) than the filtered community; yet again, this result was constrained by the reduced range in the filtered community (Fig. 3).

### Morphological traits: Wing Index

On average both communities showed similar distributions in relation to the body size of their species (WI), with slighlty higher CWM for the filtered one the first year of monitoring and higher variability over time. Other than that, the Wing Index was the most similar trait between the filtered and regional communities of all the studied traits. Similarly to the TAO, the filtered community showed a trend towards higher (but varying) multimodality (Fig. 2 and Fig. S2).

This similarity between regional and filtered communities in Wing Index is also observed in the *S*^2^ *− K* relationship, which showed no clear differences between communities (Fig. 3).

## Discussion

The variability in trait distributions between communities from a common species pool can be indicative of ecological or stochastic processes shaping the assembly and dynamics between these communities, as in urban communities and their (semi)natural source areas (Aronson *et al*., 2016; Fournier *et al*., 2020). Here we have shown that analysing different dimensions of trait distributions is necessary to understand such potential processes. Using a comprehensive dataset of butterfly communities from a regional community and a filtered urban community, we found that trait distributions from both communities are qualitatively different, but these differences are most informative when analysing a whole set of traits (morphological, behavioural, physiological) and distribution descriptors of the distributions, including their skewness, kurtosis, range, and multimodality. Furthermore, we found that trait distributions in both communities were relatively consistent across a period of six years, particularly the regional community.

Besides its statistical convenience, there is no a-priori reason to expect trait distributions to be gaussian (Gross *et al*., 2021). If anything, we might expect distributions with a relatively long tail of “rare” trait values, by analogy with Species Abundance Distributions (McGill *et al*., 2007). Trait distributions for plant species or communities are comparatively easier to obtain than for animals, and these tend to be non-gaussian (e.g. Enquist *et al*. (2017)). This has consequently pushed the development of methodologies and metrics to compare trait distributions that go beyond comparisons of Community Weighted Means (Fitzgerald *et al*., 2024; Le Bagousse-Pinguet *et al*., 2017; Maitner *et al*., 2023). We provide empirical evidence that non-gaussian trait distributions are also observed in animal communities, since in our butterfly study system none of the distributions, either from the regional or filtered communities, was unequivocally gaussian. Rather, trait distributions were skewed positively (HSI,SSI,STI,WI) or negatively (SVTI,TAO) in most cases; similarly, most distributions likewise showed positive kurtosis (Fig. 2). In addition, the DIP multimodality test showed significant departure from unimodality in all cases. In such situations, comparing distributions or inferring filtering mechanisms only from the study of community weighted means is incomplete (Gross *et al*., 2021).

The two communities studied correspond to a regional, (semi)natural community, and to a urban area potentially subject to dispersal, environmental, or stochastic filters from the regional community (Pla-Narbona *et al*., 2020; Silva *et al*., 2016; Sol *et al*., 2014). Previous research has postulated three types of environmental filtering in community assemblages resulting from non-neutral processes, namely disruptive filtering (selection of alternative optimal trait values), directional filtering (i.e., selection of optimal and reduction non-optimal values), and stabilizing filtering (i.e.reduced variance around the domminant trait values; e.g Gross *et al*. (2021); Hulme & Bernard-Verdier (2018); Loranger *et al*. (2018)), but few empirical studies have demonstrated them (e.g. (Fitzgerald *et al*., 2024; Gross *et al*., 2017)). We found, first, that trait distributions differed between our regional and filtered communities for all traits except the Wing Index (WI). In general, and despite a general higher multimodality in the filtered community, the dominant trait values coincided in both communities, whithout clear evidence of disruptive filtering. This suggests that neutral dynamics occur for dominant values (i.e., equally probable frequencies without selection (Weiher *et al*., 2011); for example, if the number of immigrants into the filtered community is proportional to the abundances in the regional community).

In contrast, the frequency of non-dominant trait values clearly differed between communities due to both reduction and exclusion of non-dominant values (e.g. in Fig. 1, extreme values found in the regional distribution are not observed in the filtered one). This, in general, subsequently led to a reduced range of the distribution in the filtered community for all traits except WI. These observations are therefore evidence of a combined filtering process, involving both directional and stabilizing responses. Directional because the reduction in trait values tends to occur in the longest tail of the asymmetric distributions; stabilizing because by filtering non-dominant values, distributions become more centered towards dominant ones. This combined filtering can only emerge in non-gaussian trait distributions: gaussian ones partly differentiate directional or stabilizing filtering based on the exlusion of one or two tails, respectively (e.g. Fig. 1 of Balazs *et al*. (2022)). Given the increasing evidence shown here and elsewhere in documenting non-gaussian trait distributions across different taxa (Enquist *et al*., 2017; Gaedke & Klauschies, 2017; Gross *et al*., 2021), reconceptualizing filtering processes in the light of such distributions is an exciting avenue for future research, particularly in animal ecology. As a final insight, the observed reduction of the range of trait values in the filtered distributions concurrently altered the statistical descriptors of the distributions. Range values are rarely evaluated when comparing communities, however our resuls clearly highlight its importance. Overall, the observed filtering and the exclusion of non-dominant species indicate deterministic selection processes in the assembly of filtered urban community that cannot be captured solely by using community-weighted means.

The observed temporal consistency of the trait distributions (Fig. 4) further indicates that the filtering processes structuring the dynamics and assembly of our study communities have a strong deterministic component. Within this overall trend, the filtered urban community showed in general higher temporal variability, which may potentially be due to a combination of factors including ecological drift and sampling errors. For example, sampling in 2020 was partially disrupted by COVID-19 lockdowns and other confinement measures, particularly in the urban community, which resulted in potential outliers in the distribution metrics (e.g. higher skweness and lower kurtosis for TAO in 2020). This higher variability in the urban communities is nevertheless expected, given the smaller community sizes (a total of 22,551 *versus* 589,316 individuals in the filterd and regional community) and species richness (54 *versus* 160 morphospecies), which potentially results in a lower capacity for portfolio effects (Schindler *et al*., 2015).

## Acknowledgements

We thank all volunteers from the CBMS (https://www.catalanbms.org/en/) and the uBMS (https://ubms.creaf.cat/en/) for providing the data needed for the study. DGC was funded by the New Zealand’s Biological Heritage National Science Challenge and by the Austrian Science Fund, with grant reference FWF ESPRIT ESP-671. The uBMS project was funded by Fundacíon Biodiversidad (2018) and the Barcelona City Hall (2019-2026). Research was supported by the projects MEDYCI Grant *PID*2020 *−* 113133*RB − I*00 funded by MCIN/ AEI /10.13039*/*501100011033 and SATURNO Grant *CNS*2022 *−* 135489 funded by MICIU/AEI*/*10.13039*/*501100011033 and by the European Union NextGenerationEU/PRTR.

## Data and code availability

The count and abundance butterfly data that support the findings of this study are available from the European Butterfly Monitor Scheme (https://butterfly-monitoring.net/), or directly from each participating Butterfly Monitoring Scheme (https://www.catalanbms.org/en and https://ubms.creaf.cat/en) via a signed license agreement. Data of species traits, their distributional metrics and R codes are available in GitHub https://github.com/garciacallejas/butterfly trait distributions.

## Author Contribution

YM and DGC conceived and designed the idea. YM and CS acquired and reviewed the data. YM and DGC analysed the data, and with CS interpreted the results. All three authors wrote the manuscript led by YM and DGC.

