## Supplementary Information for "Beyond community-weighted means: quantifying trait distributions for detecting community assembly patterns"

### Supporting Information

### Methods S1

Butterfly monitoring consists of visual identification and counts of adult butterflies during the flight season (March to the end of September in warm temperate and xeric climatic zone of the area). Counts are done weekly following the "Pollard Walk", the standard methodology in the national butterfly monitoring schemes. Monitoring of the regional community was done along 129 fixed transects in natural areas of ca. 2 km in length on average (range: 0.7 – 4.9 km). Monitoring of the filtered community was done combining fixed transects of 300 m length and random walks of fixed site-specific duration, to account for small isolated areas such as the urban ones (i.e. parks and gardens). Duration of the walk was set based on the formula:

$$T_i = (\ln(A_i + 0.75)) \times 15 \tag{E1}$$

where T is the duration in time of the free random walk (in minutes) of the site *i*, and A is the area in hectares of the site. Constant values were set to following project COCONUT (http://www.coconut.pensoft.net/pub.html), modified to reduce walking times to a maximum of 2h walk.

The duration of the schemes varied in terms of starting year, with the CBMS running since 1994 but the uBMS starting in 2018. Hence, we used data collected from 2018 for both schemes until 2023. All data collected by volunteers in both schemes is annually inspected for potential errors. Identification of individuals was done to the species level when possible ($N_{\text{CBMS}} = 569539$, $N_{\text{uBMS}} = 21565$) or, alternatively, to the genus ($N_{\text{CBMS}} = 15035$, $N_{\text{uBMS}} = 217$; which included *Agrodiaetus sp.*, *Argynnis sp.*, *Hipparchia sp.*, *Leptidea sp.*, *Limenitis sp.*, *Lysandra sp.*, *Glaucopsyche sp.*, *Gonepteryx sp.*, *Melitaea sp.*, *Melanargia sp.*, *Pieris sp.*, *Pyrgus sp.* and *Thymelicus sp.*) or family ($N_{\text{CBMS}} = 4742$, ($N_{\text{uBMS}} = 769$; included Hesperidae, Lycaenidae, Nymphalidae, Pieridae and Satyridae) level, following the nomenclature by Wiemers *et al.* (2018). Species identified to the genus or family taxa level were set to morphospecies (3.5 and 4.6% of the total observations for

CBMS and uBMS, respectively), and the values of the traits were set as the average value of the trait among the categorization (e.g. the trait value of a morphospecies set to the genus Pieris sp. was calculated as the average of the trait values of the genus).

### Methods S2

The annual abundance index calculator failed for 25 and 7 species in each community due to sporadic observations (low numbers or temporally too clumped to calculate the species phenology) within the year. To account for their contribution in the trait distribution, we extrapolated their annual abundance using linear regressions, separately for each community given species abundances differed between them. Counts were bound to a maximum of 500 individuals per species, year and community to avoid asymptotic behaviour in the abundance distributions. Both models were checked for normality and heteroskedasticity (p-value $< 0.0001$), and the latter corrected using robust standard errors from the robust covariance matrix. Analyses were done using lme4, lmtest and sandwich package in R v4.4.2 (Bates *et al.*, 2015; Zeileis, 2004; Zeileis *et al.*, 2002). Predictions provided an estimated slope of 0.015 and 0.07 for the CBMS and the uBMS respectively (p-values $<$ 10-16; Figure S1).

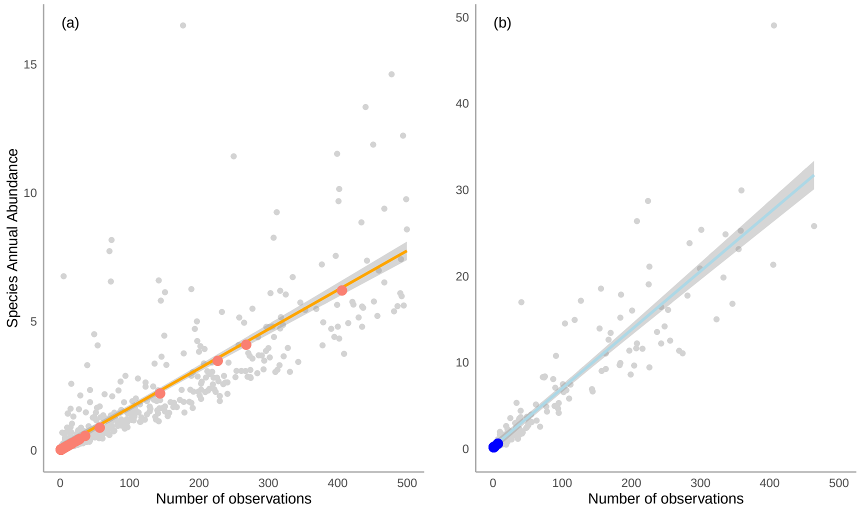


Fig. S1: Estimated annual species abundance indexes in relation to the count data for a) the regional (CBMS), and b) the filtered community pool (uBMS). Lines relate to the estimated model slopw and 95% interval confidence. Colour dots relate to predicted abundances from missing failes annual indexes calculations

.

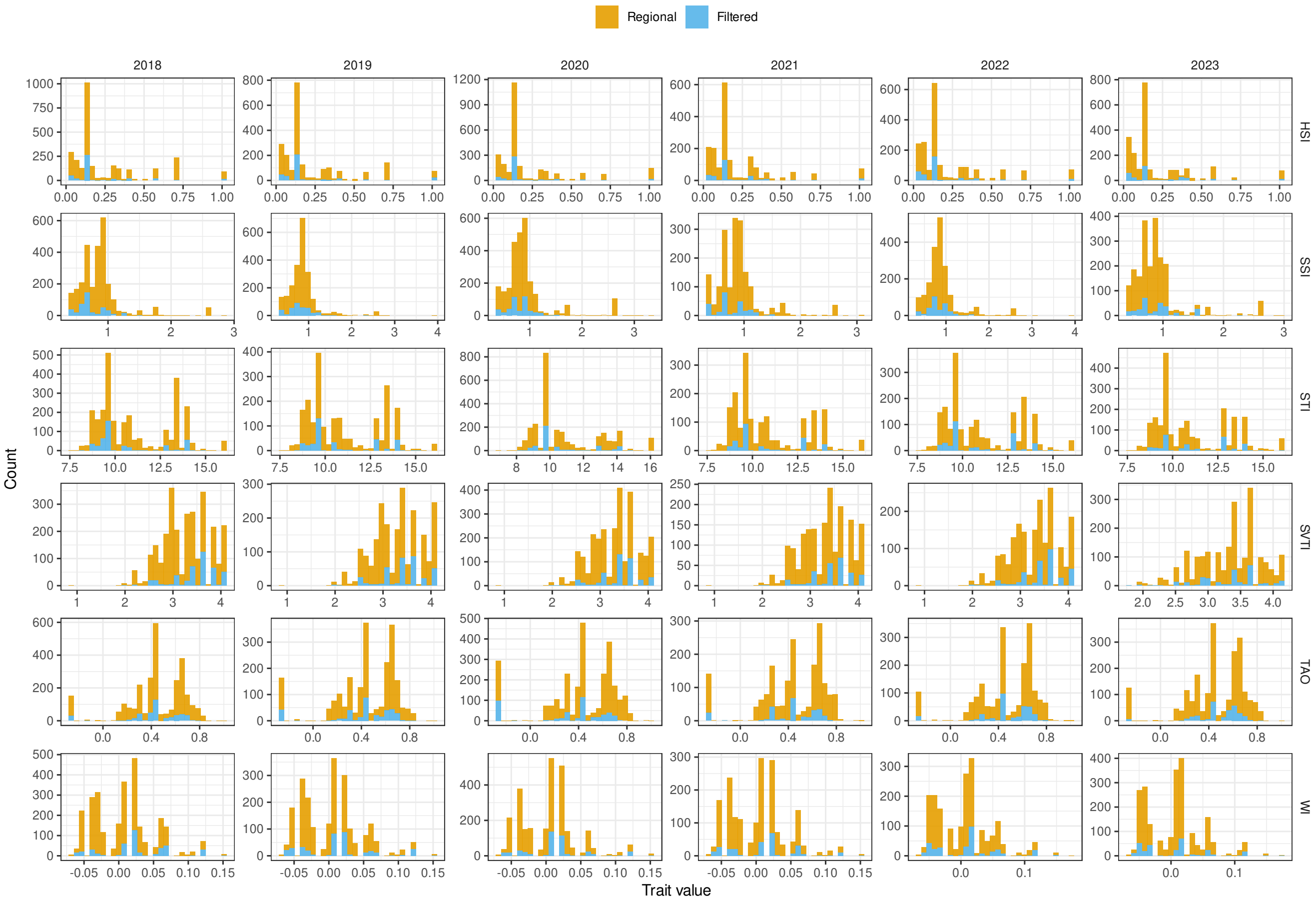


Fig. S2: Distributions of six traits from a regional community (natural areas surrounding the municipality of Barcelona) and a filtered community (the municipality of Barcelona) from 2018 to 2023. For the definition of the traits used, see Methods of the main text.
